# The sowing date changed the temperature and light conditions in the field modified the cadmium content of brown rice (*Oryza sativa* L.) by regulating the expression of Cd-related genes

**DOI:** 10.1101/2022.01.24.477481

**Authors:** Zhongwen Rang, Huan Xiao, Zhenxie Yi, Xuehua Wang, Qiaomao Chen, Yu Kuang, Hejun Ao

## Abstract

Cadmium (Cd) contamination in rice is a potential health hazard when ingested through the food chain worldwide. Reducing the Cd content in rice through agronomic measures is an effective way to reduce the risk of Cd contamination to human health. In order to clarify the correlation between temperature and light conditions and Cd accumulation (Cd-A) and Cd content of brown rice (CdBR) during the field growth period (FGP) of rice, consequently provide a theoretical basis for the selection of sowing date (SD) for “Low-Cd-Rice” production, field experiment with different SDs was carried out by using two rice varieties with different Cd accumulation characteristics (Luliangyou 996, V1, a high Cd accumulation variety; Zhuliangyou 819, V2, a low Cd accumulation variety). The results showed that the temperature and light factors such as mean soil temperature (ST), mean air temperature (AT), soil accumulation temperature (SAT), air accumulation temperature (AAT), ultraviolet radiation accumulation (UR), photosynthetic radiation accumulation (PR), light intensity accumulation (I) and sunshine hours accumulation (SH) varied to different degrees under different SDs; The difference in CdBR in two varieties could be up to 2.82 and 8.48 times respectively among SDs, with the CdBR of S4 and S5 of V2 being lower than the national standard of 0.2 mg/kg. The relative expression of *OsIRT1* in the root system was significantly positively correlated with ST, SAT, AT, AAT, and SH, while *OsNramp5, OsNramp1*, and *OsHMA3* showed significant negative correlations with ST, SAT, AT, AAT, and SH in relative expression in the root system; *OsIRT1* expressed in the roots of V1 was significantly negatively correlated with CdBR, while *OsHMA3* expression was significantly positively correlated with *CdBR; OsLCD, OsNramp1*, and *OsHMA3* expression in the roots of V2 were significantly positively correlated with Cd-A and CdBR, while *OsIRT1* in the roots of V2 and *OsLCT1* in the leaves were significantly negatively correlated with Cd-A; The expression of *OsNramp5* in roots was significantly negatively correlated with Cd-A and CdBR in both V1 and V2. Bias correlation analysis showed that ST, SAT, AT, and AAT were significantly negatively correlated with both Cd-A and CdBR; SH was significantly negatively correlated with CdBR in V1. Summarily, the temperature and light conditions during the FGP of rice and their regulation of the expression levels of related genes could be changed by sowing selection, so as to achieve safe production of rice under Cd-contaminated fields.

## 1 Introduction

Rice is one of the three major food crops in the world and more than 50% of the world’s population as the staple food. In China, rice production and consumption play a leading role in food production. More than 65% of the population in China takes rice as the staple food, which is the cornerstone of China’s food security. With the rapid development of modern chemical agriculture, global cadmium (Cd) pollution is becoming more and more serious[1]. A large amount of Cd enters to the farmland ecosystem, resulting in excessive Cd content in farmland soil. In recent years, the problem of heavy metal pollution of paddy soil and rice in China has become increasingly serious[2], among which Cd pollution is the most serious. It was studied that approximately 1.3 × 10^4^ ha of cultivated soil was contaminated by Cd and about 5.0 × 10^4^t of Cd contaminated rice was produced every year[3]. The incidents of “cadmium rice” and “toxic rice” have occurred frequently, which has attracted extensive attention in the society[4]. Cd has strong biological mobility and is easily absorbed and accumulated by plants, which can be ingested into human body through soil-crop-food chain system, becoming the main source of Cd intake and causing potential harm to human health[5]. Therefore, reducing the content of Cd in rice has become one of the effective ways to reduce the harm of Cd pollution to human health[6].

Cd is a non-essential element for rice growth, but it can be absorbed from the soil through the transport channel of essential mineral nutrient elements in rice roots[7], realize the distribution between aboveground stems and leaves through xylem loading and transportation, and further migrate to grains through phloem, and finally complete the accumulation in grains[8,9]. The absorption, transport and distribution of Cd by rice are related to varieties[10], soil types and pH [11,12], irrigation methods[13,14], cultivation modes[15], etc. Positive progress has also been made in the research on the mechanisms of cadmium tolerance and the molecular mechanisms of Cd absorption and accumulation in rice[3, 16–18]. For example, *OsABCG36* enhances the tolerance of rice to Cd by expelling Cd from root cells[19]; The transporter *OsHMA3* can transport Cd into vacuoles, reducing the transport of Cd to aboveground, reducing the toxic effect of Cd[20]; Cd enters the vessels through transporters, and then transfers upward by transpiration and root pressure is the key to determine the accumulation of Cd in rice shoot and grain[21]; The transfer of Cd from vascular to DVB (Diffuse Vascular Bundles) is the key to the accumulation of Cd into grains. Selecting varieties with small DVB area in the first stem node is conducive to reducing the accumulation of Cd in brown rice[22]. Not exclusive to these studies have laid a good theoretical and technical foundation for the cultivation of rice varieties with high yield, high quality and low Cd accumulation, and the cultivation regulation of Cd absorption and accumulation in rice as well.

Temperature and light are two particularly important environmental factors for plant survival, growth and development[23]. Different temperature and light conditions affect rice growth period and material accumulation, thus affecting rice yield composition[24], grain quality[25], and the differential absorption and accumulation of Cd by rice[26]. Differences in Cd content of rice under different yield levels was also proved[27]. Different genotypes and in the same rice variety in different seasons and locations which considered an environmentally variable variety, suggested genotypic differences in Cd uptake and accumulation and possible gene-environment interactions[26]. It was suggested that temperature was the main factor caused the difference in Cd content of these environmentally variable varieties[28]. In addition, the Cd absorption of rice was most sensitive to the temperature changes in tillering and grain filling stage, low temperature in the early growth stage and high temperature in the late growth stage could promote the accumulation of cadmium in rice grains[29], that’s the reason why the content of Cd in grains and rachis of late rice is higher than that of early rice. However, the physiological mechanisms by which temperature affects Cd uptake and transport in rice were seldom reported. Additionally, light conditions affect the growth and development of rice and inevitably affect various physiological metabolic processes of Cd uptake, translocation and accumulation in rice under Cd contaminated conditions[30,31], although details of the ways in which light affects the uptake of Cd in rice and the related physiological metabolic mechanisms have not been reported.

The present work was carried out aiming to elucidate the uptake and accumulation of Cd in different types of rice varieties under different light and temperature conditions by setting different SDs, and the interrelationship between Cd uptake and accumulation and the expression of Cd uptake and translocation-related genes, so as to deepen the physiological mechanism of Cd uptake and accumulation in rice under temperature and light conditions, and provide a theoretical basis for the selection of rice varieties and their suitable sowing seasons in Cd-polluted areas of China, the determination of rice planting systems, and the effective regulation of Cd content in rice.

## 2 Materials and Methods

### 2.1. Experimental varieties and field

Two-line Early Hybrid Rice Luliangyou 996 (V1) and Zhuliangyou 819 (V2) were used in this study. V1 is the main variety of double cropping early rice in the middle and lower reaches of the Yangtze River, with an average growth period of 109.7 days (from sowing to harvesting) and characteristics of relative high Cd accumulation. V2 is an emergency early rice variety with low Cd accumulation popularized in Hunan Province (light incidence areas of rice blast) with an average growth period of about 106.0 days. Field experiment was conducted in 2018 in Yonghe village, Yanxi Town, Liuyang City of Hunan Province (Comprehensive Teaching and Experimental Base of Hunan Agricultural University), the cadmium content (total cadmium) of experimental field soil was 0.47 ± 0.07mg/kg, and soil pH was 5.4 ± 0.3. The experimental soil type was loam, and the basic soil nutrients were as follows: organic matter, 22.71g/kg; total nitrogen, 1.63 g/kg; total phosphorus, 1.56g/kg; alkali hydrolyzable nitrogen, 133.51 mg/kg; available phosphorus, 38.59mg/kg; and available potassium, 134.26mg/kg.

### 2.2. Experimental design

Every 15 days from April 22, 2018, to July 6, 2018, totally six different SD treatments were set by strip-plot design with 2 repetitions in each variety, 24 plots were set with an area of 40m^2^(4m × 10m) each. Separate water inlet and drainage ditches were set on both sides of every plot, and ridges were made among treatments covered with plastic film, the variety interval was 0.8m and the repetition interval was 0.4m.

### 2.3. Cultivation and field management

Exactly two of 25 days old seedlings were transplanted with specification of 20cm×20cm in each transplanting point. The fertilization program was as follows: the proportion of base fertilizer, applied 2 days before transplanting, and tiller fertilizer, applied 20 days after transplanting, was 6:4. 180kg/hm^2^ pure nitrogen (urea, nitrogen content is 46.4%), 90kg/hm^2^ P_2_O_5_ (calcium superphosphate, P_2_O_5_ content is 12%, as base fertilizer), and 145kg/hm^2^ K_2_O (potassium chloride, K_2_O content is 60%, as the base fertilizer: top-dressing fertilizer = 0.5:0.5) were applied in each treatment plot. Plant protection measures was uniformly managed according to local regulations, and there were no obvious diseases, pests, weeds and meteorological disasters during experimental period.

### 2.4. Soil and meteorological data acquisition

The meteorological data were recorded once an hour by using the micro meteorological station (Vantage Pro 2, USA) installed in the field. The main meteorological indicators include air temperature (AT, °C), soil temperature (ST, °C), ultraviolet radiation (UR, MJ), photosynthetic radiation (PR, KW/m^2^), light intensity (I, Klux), sunshine hours (SH, h), soil pH, air CO_2_ concentration (CO2, ppm), atmospheric pressure (AP, hpa), rainfall (RF, mm), etc.

### 2.5. Determination of Cd content in brown rice and Cd accumulation in plant

Processing grains (each sample) into brown rice, and screened through a 100-mesh sieve after crushed with a stainless-steel crusher. Concentrated nitric acid and perchloric acid (V nitric acid: V Perchloric acid = 4:1) were used to wet digestion. The cadmium content of crushed samples was determined by atomic spectrophotometer (Graphite Furnace). Dry weight (DW, kg) and Cd Content (mg/kg) of root, leaf and stem, and spike of each variety were also determined for computing the Cd accumulation (Cd-A) by plant in the experimental condition with the formula as follows:

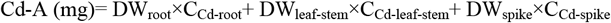

### 2.6. Real-time PCR analysis

Root and leaf samples were taken by liquid nitrogen at rice maturity and then stored in a –80°C refrigerator. Genes related to Cd uptake and transport e.g., *OsLCD, OsIRT1, OsIRT2, OsLCT1, OsNramp1, OsNramp5* and *OsHMA3*, expressed in root and genes *OsLCD, OsIRT1, OsIRT2* and *OsLCT1* expressed in leaf were analyzed by using Real-time RT-PCR (the primer sequences used for qRT-PCR are shown in Table 1., Synthesis by Invitrogen, Beijing). The total RNA samples were isolated by TRIzol reagent (TIANGEN BIOTECH, Beijing). Then the RNA purity and concentration was measured by using the NanoPhotometer spectrophotometer (IMPLEN, CA, USA). After detecting, cDNA was synthesized using 2 μg RNA using the PrimeScript™ RT reagent Kit with gDNA Eraser (TaKaRa). Gene specific primers for quantitative real-time PCR (qRT-PCR) analysis were designed using Primer 5.0 by Allwegene Technology (Allwegene Technology Co., Ltd. Beijing, China). The *ACTIN* gene was used as internal reference gene. qRT-PCR reaction was performed using SYBR^®^ Premix Ex Taq™ II (Tli RNaseH Plus) and was conducted on ABI 7500 Real-time Detection System (Thermo Fisher Scientific, USA). The PCR reaction was carried out with the following reaction conditions: 95°C for 30s; followed by 45 cycles of 95°C for 5s, 60°C for 40s. Samples for qRT-PCR were run in 3 biological replicates with 3 technical replicates and the data were represented as the mean ± SD (n = 3) for Student’s t-test analysis. The relative gene expression was calculated using the 2^-ΔΔCT^ algorithm[32].

**Table 1.**
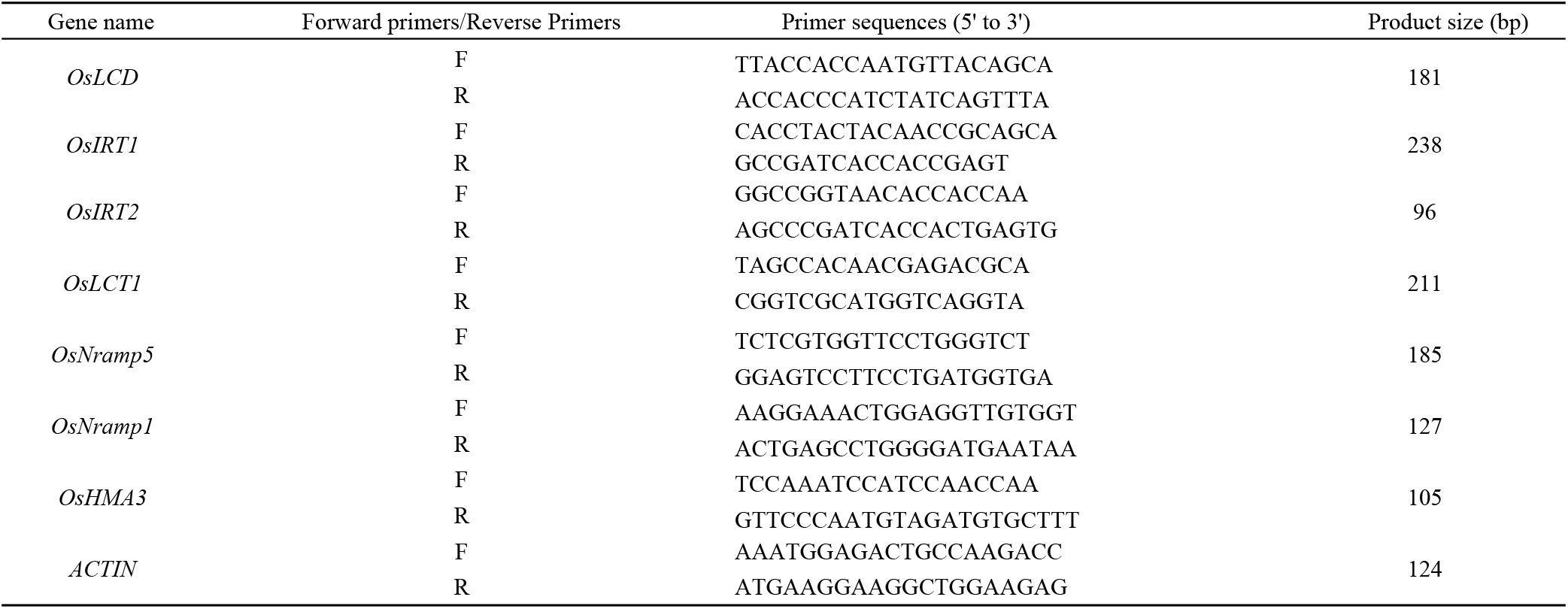
Target gene primer sequences for Real Time PCR analysis.

### 2.7. Statistical analysis

Excel 2010 was used for data processing and plotting, and SPSS 22.0 was used for descriptive statistics, independent sample t-test, single sample t-test, ANOVA, multiple comparison, Pearson correlation analysis, Partial correlation analysis, etc.

## 3 Results

### 3.1 Variations of temperature and light indicators under different SD treatments

Independent sample t-test results shown that all of average values of temperature and light indicators monitored from two varieties shown no significant difference (Table 2.), indicated that the changes in the indicators of the two experimental varieties were consistent in response to the SD treatments. The differences of average temperatures (ST, AT), accumulative temperatures (SAT, AAT), and light factors (UR, PR, I, SH) among different SD treatments were due to the differences of stage and length of the FGP of each treatment. Temperature factors e.g., ST, AST, AT and AAT showed less variation among treatments with CVs of 4.23%-5.50% and 4.10%-5.95% for the two varieties respectively, while light factors e.g., UR, PR and I showed more variation among treatments with CVs of 14.08%-16.75% and 13.66%-16.80% for the two varieties respectively. In addition, the SH varied less across treatments, with an average CV of 6.58% for two varieties.

**Table 2.** Characterization of indicators related to temperature and light factors under different SD treatments. V, indicates variety, V1, Lu Liangyou996, V2, Zhu Liangyou 819; T, indicates different sowing treatments; S1–S6 indicates the different treatments in the specific sowing date (SD); ST, soil daily average temperature during field growth period(FGP); AST, soil accumulated temperate; AT, daily average air temperature; AAT, accumulated air temperature; UR, accumulative of daily ultraviolet radiation in FGP; PR, accumulative daily photosynthetic radiation in FGP; I, accumulative daily illuminance in FGP; SH, accumulative sunshine hours in FGP; St. D means the standard deviation; CV (%) indicates the Coefficient of variation; Cohen’s d indicates the T–value of the independent sample t–test, and Sig. indicates the two–tail significance at 5% level.

| V | Statistical Value | T | ST /°C | AST /°C | AT /°C | AAT /°C | UR /MJ | PR kW/m² | I /Klux | SH /h |
| --- | --- | --- | --- | --- | --- | --- | --- | --- | --- | --- |
| V1 |  | S1 | 28.16 | 2365.20 | 26.98 | 2293.37 | 8.00 | 1312.35 | 4461.53 | 776.70 |
|  |  | S2 | 28.47 | 2362.91 | 27.46 | 2306.58 | 8.78 | 1402.33 | 4605.51 | 778.50 |
|  |  | S3 | 28.97 | 2404.63 | 28.14 | 2363.79 | 10.18 | 1562.60 | 5181.88 | 777.60 |
|  |  | S4 | 28.72 | 2383.40 | 27.82 | 2337.18 | 11.88 | 1767.25 | 6147.00 | 752.30 |
|  |  | S5 | 27.70 | 2299.03 | 26.54 | 2229.09 | 12.40 | 1898.61 | 6679.64 | 728.70 |
|  |  | S6 | 25.72 | 2109.14 | 24.37 | 2022.65 | 11.13 | 1736.59 | 6160.88 | 655.80 |
|  | Mean |  | 27.96 | 2320.72 | 26.89 | 2258.78 | 10.40 | 1613.29 | 5539.41 | 744.93 |
|  | St. D |  | 1.18 | 109.51 | 1.36 | 124.33 | 1.74 | 227.14 | 918.40 | 47.89 |
|  | CV (%) |  | 4.23% | 4.72% | 5.05% | 5.50% | 16.75% | 14.08% | 16.58% | 6.43% |
| V2 |  | S1 | 28.17 | 2394.23 | 27.00 | 2322.23 | 8.10 | 1326.26 | 4458.52 | 783.70 |
|  |  | S2 | 28.47 | 2391.68 | 27.47 | 2334.63 | 8.85 | 1415.11 | 4590.30 | 786.30 |
|  |  | S3 | 28.97 | 2404.63 | 28.14 | 2363.79 | 10.18 | 1562.60 | 5181.88 | 777.60 |
|  |  | S4 | 28.72 | 2383.40 | 27.82 | 2337.18 | 11.88 | 1767.25 | 6147.00 | 752.30 |
|  |  | S5 | 27.70 | 2299.03 | 26.54 | 2229.09 | 12.40 | 1898.61 | 6679.64 | 728.70 |
|  |  | S6 | 25.81 | 2090.67 | 24.47 | 2006.95 | 11.13 | 1734.90 | 6229.87 | 655.20 |
|  | Mean |  | 27.97 | 2327.27 | 26.91 | 2265.65 | 10.42 | 1617.46 | 5547.87 | 747.30 |
|  | St. D |  | 1.15 | 122.10 | 1.32 | 134.89 | 1.70 | 220.92 | 931.94 | 50.23 |
|  | CV (%) |  | 4.10% | 5.25% | 4.92% | 5.95% | 16.32% | 13.66% | 16.80% | 6.72% |
|  | Cohen's d |  | -0.028 | -0.092 | -0.025 | -0.098 | -0.029 | -0.032 | -0.016 | -0.084 |
|  | Sig. |  | 0.978 | 0.929 | 0.981 | 0.924 | 0.978 | 0.975 | 0.988 | 0.935 |

### 3.2 Relative expression of genes related to Cd uptake and transport under different SDs

As shown in Table 3., with the exception of *OsIRT2*, the relative expression variation of genes related to Cd uptake and translocation in roots and leaves at maturity stage under the SDs was consistent in the two rice varieties. The genes showing high variation(CV>36%) in relative expression in the root system under different SD treatments in both varieties were *OsIRT1*, *OsLCT1*, *OsNramp5* and *OsHMA3; OsIRT2* showed small variation in expression in both varieties (CV<15%); while *OsLCD* showed high variation in expression in V1 and moderate variation in V2 (16%<CV<35%) and *OsNramp1* was moderately variable in V1 and highly variable in V2. In the leaves, *OsIRT1* showed high variation in both varieties, *OsLCD* and *OsIRT2* showed small variation in both varieties, but the relative expression of *OsLCT1* differed between the two varieties due to differences SD treatments, with high variation in V1 and moderate variation in V2.

**Table 3.**
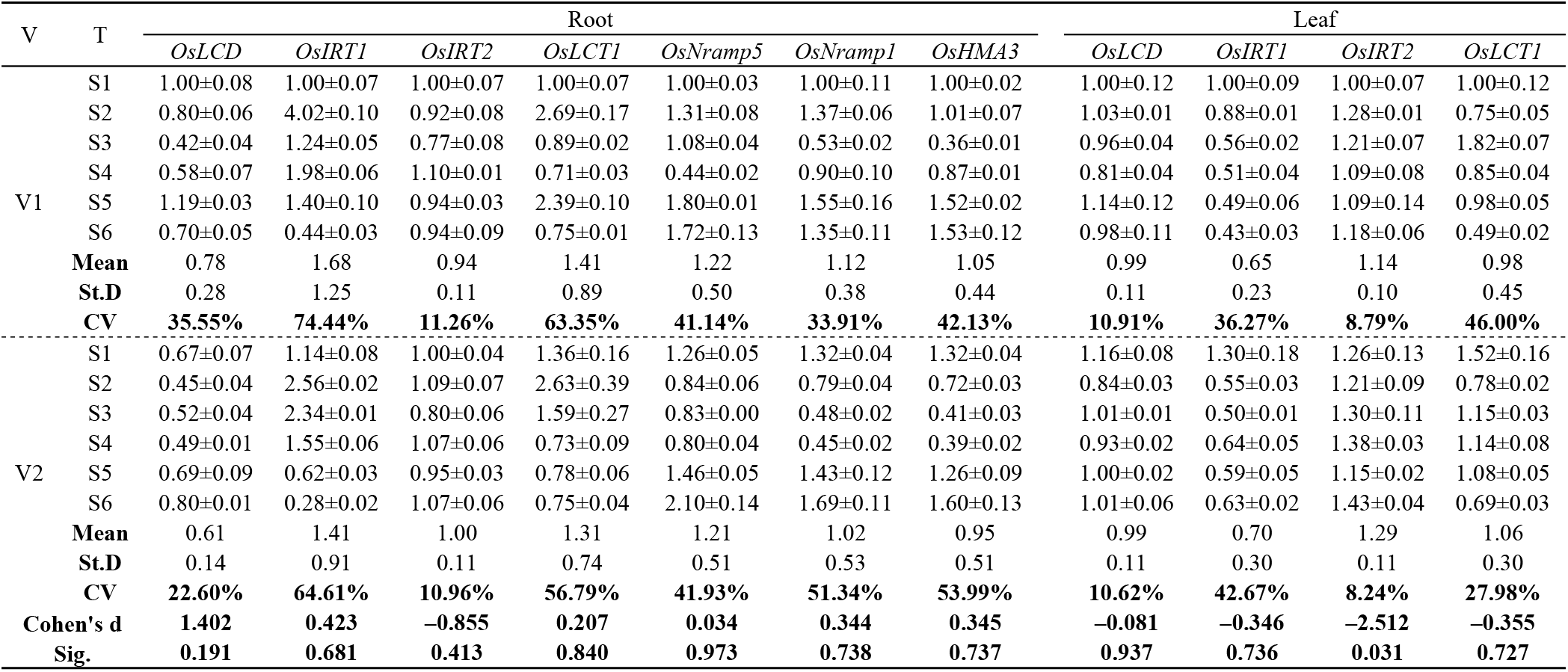
Relative expression of genes related to Cd uptake and transport at maturity stage in two rice varieties under different SD treatments.

| V | T | Root |  |  |  |  |  |  | Leaf |  |  |  |
| --- | --- | --- | --- | --- | --- | --- | --- | --- | --- | --- | --- | --- |
|  |  | *OsLCD* | *OsIRT1* | *OsIRT2* | *OsLCT1* | *OsNramp5* | *OsNramp1* | *OsHMA3* | *OsLCD* | *OsIRT1* | *OsIRT2* | *OsLCT1* |
| V1 | S1 | 1.00±0.08 | 1.00±0.07 | 1.00±0.07 | 1.00±0.07 | 1.00±0.03 | 1.00±0.11 | 1.00±0.02 | 1.00±0.12 | 1.00±0.09 | 1.00±0.07 | 1.00±0.12 |
|  | S2 | 0.80±0.06 | 4.02±0.10 | 0.92±0.08 | 2.69±0.17 | 1.31±0.08 | 1.37±0.06 | 1.01±0.07 | 1.03±0.01 | 0.88±0.01 | 1.28±0.01 | 0.75±0.05 |
|  | S3 | 0.42±0.04 | 1.24±0.05 | 0.77±0.08 | 0.89±0.02 | 1.08±0.04 | 0.53±0.02 | 0.36±0.01 | 0.96±0.04 | 0.56±0.02 | 1.21±0.07 | 1.82±0.07 |
|  | S4 | 0.58±0.07 | 1.98±0.06 | 1.10±0.01 | 0.71±0.03 | 0.44±0.02 | 0.90±0.10 | 0.87±0.01 | 0.81±0.04 | 0.51±0.04 | 1.09±0.08 | 0.85±0.04 |
|  | S5 | 1.19±0.03 | 1.40±0.10 | 0.94±0.03 | 2.39±0.10 | 1.80±0.01 | 1.55±0.16 | 1.52±0.02 | 1.14±0.12 | 0.49±0.06 | 1.09±0.14 | 0.98±0.05 |
|  | S6 | 0.70±0.05 | 0.44±0.03 | 0.94±0.09 | 0.75±0.01 | 1.72±0.13 | 1.35±0.11 | 1.53±0.12 | 0.98±0.11 | 0.43±0.03 | 1.18±0.06 | 0.49±0.02 |
|  | Mean | 0.78 | 1.68 | 0.94 | 1.41 | 1.22 | 1.12 | 1.05 | 0.99 | 0.65 | 1.14 | 0.98 |
|  | St.D | 0.28 | 1.25 | 0.11 | 0.89 | 0.50 | 0.38 | 0.44 | 0.11 | 0.23 | 0.10 | 0.45 |
|  | CV | 35.55% | 74.44% | 11.26% | 63.35% | 41.14% | 33.91% | 42.13% | 10.91% | 36.27% | 8.79% | 46.00% |
| V2 | S1 | 0.67±0.07 | 1.14±0.08 | 1.00±0.04 | 1.36±0.16 | 1.26±0.05 | 1.32±0.04 | 1.32±0.04 | 1.16±0.08 | 1.30±0.18 | 1.26±0.13 | 1.52±0.16 |
|  | S2 | 0.45±0.04 | 2.56±0.02 | 1.09±0.07 | 2.63±0.39 | 0.84±0.06 | 0.79±0.04 | 0.72±0.03 | 0.84±0.03 | 0.55±0.03 | 1.21±0.09 | 0.78±0.02 |
|  | S3 | 0.52±0.04 | 2.34±0.01 | 0.80±0.06 | 1.59±0.27 | 0.83±0.00 | 0.48±0.02 | 0.41±0.03 | 1.01±0.01 | 0.50±0.01 | 1.30±0.11 | 1.15±0.03 |
|  | S4 | 0.49±0.01 | 1.55±0.06 | 1.07±0.06 | 0.73±0.09 | 0.80±0.04 | 0.45±0.02 | 0.39±0.02 | 0.93±0.02 | 0.64±0.05 | 1.38±0.03 | 1.14±0.08 |
|  | S5 | 0.69±0.09 | 0.62±0.03 | 0.95±0.03 | 0.78±0.06 | 1.46±0.05 | 1.43±0.12 | 1.26±0.09 | 1.00±0.02 | 0.59±0.05 | 1.15±0.02 | 1.08±0.05 |
|  | S6 | 0.80±0.01 | 0.28±0.02 | 1.07±0.06 | 0.75±0.04 | 2.10±0.14 | 1.69±0.11 | 1.60±0.13 | 1.01±0.06 | 0.63±0.02 | 1.43±0.04 | 0.69±0.03 |
|  | Mean | 0.61 | 1.41 | 1.00 | 1.31 | 1.21 | 1.02 | 0.95 | 0.99 | 0.70 | 1.29 | 1.06 |
|  | St.D | 0.14 | 0.91 | 0.11 | 0.74 | 0.51 | 0.53 | 0.51 | 0.11 | 0.30 | 0.11 | 0.30 |
|  | CV | 22.60% | 64.61% | 10.96% | 56.79% | 41.93% | 51.34% | 53.99% | 10.62% | 42.67% | 8.24% | 27.98% |
| Cohen's d |  | 1.402 | 0.423 | -0.855 | 0.207 | 0.034 | 0.344 | 0.345 | -0.081 | -0.346 | -2.512 | -0.355 |
| Sig. |  | 0.191 | 0.681 | 0.413 | 0.840 | 0.973 | 0.738 | 0.737 | 0.937 | 0.736 | 0.031 | 0.727 |

Two-way ANOVA showed (Table 4. and Table 5.) that at maturity stage, the genes whose relative expression did not differ among rice varieties but differed significantly among SD treatments were *OsLCT1* and *OsNramp5* in the roots and *OsLCD* in the leaves, respectively; while the genes whose relative expression differed significantly among varieties and among SD treatments were *OsLCD, OsIRT1, OsIRT2, OsNramp1* and *OsHMA3* in the roots, and *OsIRT1* and *OsIRT2* and *OsLCT1* in the leaves, respectively.

**Table 4.** Two–way ANOVA for relative expression of genes related to Cd uptake and transport in the roots of two rice varieties at maturity stage under different SD treatments

| Sources of error/genes | *OsLCD* |  | *OsIRT1* |  | *OsIRT2* |  | *OsLCT1* |  | *OsNramp5* |  | *OsNramp1* |  | *OsHMA3* |  |
| --- | --- | --- | --- | --- | --- | --- | --- | --- | --- | --- | --- | --- | --- | --- |
|  | F | P | F | P | F | P | F | P | F | P | F | P | F | P |
| V | 114.35 | 0.00 | 142.82 | 0.00 | 6.10 | 0.02 | 3.44 | **0.08** | 0.20 | **0.66** | 12.26 | 0.02 | 20.36 | 0.00 |
| T | 78.36 | 0.00 | 1337.52 | 0.00 | 14.78 | 0.00 | 139.81 | **0.00** | 295.24 | **0.00** | 166.20 | 0.00 | 313.15 | 0.00 |
| V×T | 37.67 | 0.00 | 248.68 | 0.00 | 2.24 | 0.08 | 42.47 | 0.00 | 49.74 | 0.00 | 34.55 | 0.00 | 32.68 | 0.00 |

**Table 5.**
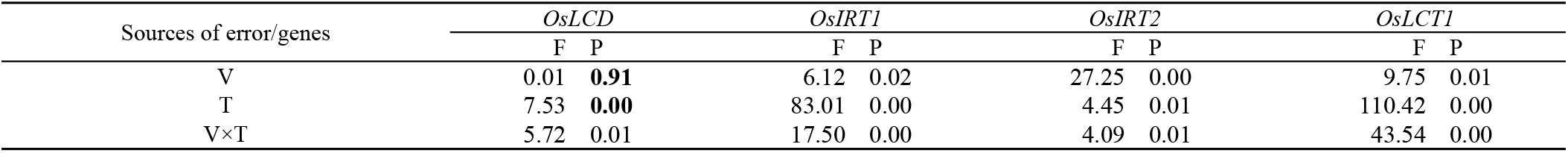
Two–way ANOVA of leaf relative expression of genes related to Cd transport at maturity stage in two rice varieties under different sowing treatments

### 3.3 CdBR under different SDs

As shown in figure 1., average CdBR in Zhu Liangyou 819 (V2) was 0.372 mg/kg, higher than Lu Liangyou 996 (V1) of 0.333 mg/kg by 11.71%, but there was no significant difference in cadmium content of brown rice (CdBR) between two varieties under different SDs (*p*=0.278). CdBR in two varieties presented significant differences (*p* < 0.05) and high variances with the CV of 47.26% in V1 and 73.99% in V2 in different SD treatments, respectively. Similarly, S6 treatment was get more higher CdBR than other treatments in two varieties, while S4s were of the lowest; CdBR in S6 were 2.82 and 8.48 times to S4 in V1 and V2, respectively. Interestingly, CdBR of S1 to S5 were lower than the criteria of CXS 193-1995 (0.4 mg/kg) [33] in V1, and S2 to S5 in V2. It was noteworthy that CdBR in S4 and S5 in V2 lower than the national standard of 0.2 mg/kg (Chinese National Standard GB 2762—2012).

**Figure 1.**
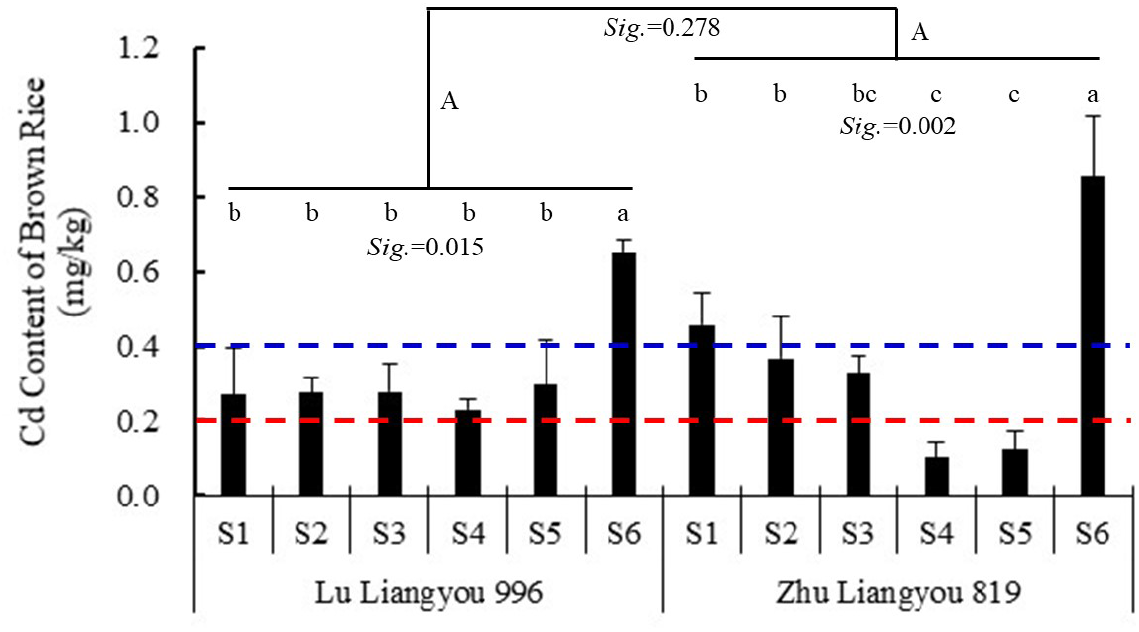
Cd content of brown rice in different sowing date treatments in Luliangyou 996 and Zhu Liangyou 819. Bars indicate ±St.D; Capital letters indicate significance between varieties and lowercase letters among different treatments at the level of 5% by using LSD. The blue and red dotted lines indicate the national standard (0.2mg/kg) and the international limit standard (0.4mg/kg) of cadmium content in brown rice, respectively.

### 3.4 Correlation between temperature and light factors and relative expression of genes related to Cd uptake and transport

Table 3. showed that there were inter-varietal differences in the expression of *OsIRT2* in leaves under different SD treatments, hence Pearson correlation analyses between the relative expression of *OsIRT2*-leaf in each variety with temperature and light factors, respectively. However, the relative expression of *OsIRT2* in the leaves of both varieties was significantly correlated with neither temperature nor light factors. Gene relative expressions in SDs with no inter-varietal differences were variety-integrated analyzed by Pearson correlation (Tabel 6.).The relative expression of *OsIRT1* in the root system was significantly positively correlated to ST, SAT, AT, AAT and SH, whereas the relative expression of *OsNramp5*, *OsNramp1* and *OsHMA3* in root were significantly negatively correlated with ST, SAT, AT, AAT and SH. Additionally, expression of *OsIRT1* in leaf was significantly negatively correlated to UR, PR, and I, while expression of *OsLCT1* in leaf was significantly positively correlated with ST, SAT, AT, and AAT.

### 3.5 Correlations between relative genes expression and Cd-A and Cd-BR

The correlations between Cd uptake and transport genes expression, Cd-A and CdBR were analyzed in groups according to whether there were varietal differences in gene expression in response to SDs (Table 7). Group 1 analyzed separately for those with varietal differences in gene expression in response to SDs, and Group 2 analyzed two rice varieties together for those without varietal differences. *OsIRT1* expressed in root of V1 was negatively significantly correlated to CdBR, while a positively significantly correlation between the expression of *OsHMA3* with CdBR; There were statistically significant correlations between the other genes expression in root or/and leaf neither with Cd-A, nor with CdBR. *OsLCD, OsNramp1*, and *OsHMA3* expressed in root of V2 were positively significantly correlated to Cd-A and CdBR, while there were negatively significant correlations between *OsIRT1* in root and *OsLCT1* in leaf of V2 only with Cd-A. Additionally in Group 2, only the expression of *OsNramp5* in root shown negatively correlations with Cd-A and CdBR.

### 3.6 Partial correlations between temperature and light factors and Cd-A and CdBR

Based on the results in Tables 6. and Table 7., the correlations between temperature and light factors and Cd-A and CdBR were further analyzed by using partial correlation analysis (Table 8.). ST, SAT, AT, AAT, and SH were significantly negatively correlated to CdBR by positively regulating the expressions of *OsIRT1* and negatively regulating *OsHMA3* in root of V1; although UR, PR, and I significantly negatively correlated to the expression of *OsIRT1* in leaf of V1, there were statistically relationships between UR, PR, and I neither with Cd-A, nor with CdBR, respectively. Temperature factors were significant affected Cd-A but not CdBR in V2, nevertheless none by light factors either in Cd-A or in CdBR. ST, SAT, AT, and AAT were negatively correlated to Cd-A in V2 by positively regulating the expression of *OsIRT1* in root and *OsLCT1* in leaf, while negatively correlated to Cd-A by negatively regulating of *OsNramp1* and *OsHMA3* in root. Additionally, ST was significantly negatively correlated with CdBR both in V1 and V2 by negatively regulating the expression of *OsNramp5* in root.

**Table 6.** Correlation between temperature and light factors and relative expression of genes related to Cd uptake and transport. “–Leaf” means the expression of genes in the leaf; * indicate significant difference at the level of 0.05%, and ** at 0.01%.

| Genes | ST | SAT | AT | AAT | UR | PR | I | SH |
| --- | --- | --- | --- | --- | --- | --- | --- | --- |
| *OsLCD* | –0.387 | –0.341 | –0.422 | –0.383 | 0.045 | 0.119 | 0.187 | –0.276 |
| *OsIRT1* | **0.613\*** | **0.576\*** | **0.617\*** | **0.591\*** | –0.350 | –0.401 | –0.482 | **0.627\*** |
| *OsIRT2* | –0.224 | –0.179 | –0.237 | –0.198 | 0.064 | 0.068 | 0.127 | –0.197 |
| *OsLCT1* | 0.273 | 0.305 | 0.258 | 0.289 | –0.360 | –0.342 | –0.400 | 0.423 |
| *OsNramp5* | **–0.821\*\*** | **–0.807\*\*** | **–0.828\*\*** | **–0.824\*\*** | 0.213 | 0.312 | 0.379 | **–0.726\*\*** |
| *OsNramp1* | **–0.724\*\*** | **–0.670\*** | **–0.750\*\*** | **–0.708\*** | 0.122 | 0.221 | 0.301 | **–0.582\*** |
| *OsHMA3* | **–0.814\*\*** | **–0.739\*\*** | **–0.839\*\*** | **–0.780\*\*** | 0.149 | 0.250 | 0.344 | **–0.669\*** |
| *OsLCD*–leaf | –0.212 | –0.139 | –0.243 | –0.180 | –0.140 | –0.076 | –0.036 | –0.064 |
| *OsIRT1*–leaf | 0.155 | 0.286 | 0.109 | 0.229 | **–0.717\*\*** | **–0.700\*** | **–0.641\*** | 0.414 |
| *OsLCT1*–leaf | **0.588\*** | **0.605\*** | **0.582\*** | **0.604\*** | –0.175 | –0.218 | –0.263 | 0.563 |
| *OsIRT2*–Leaf–V1 | –0.030 | –0.119 | 0.007 | –0.072 | –0.060 | –0.070 | –0.143 | –0.051 |
| *OsIRT2*–Leaf–V2 | –0.388 | –0.476 | –0.351 | –0.437 | 0.155 | 0.120 | 0.189 | –0.520 |

**Table 7.**
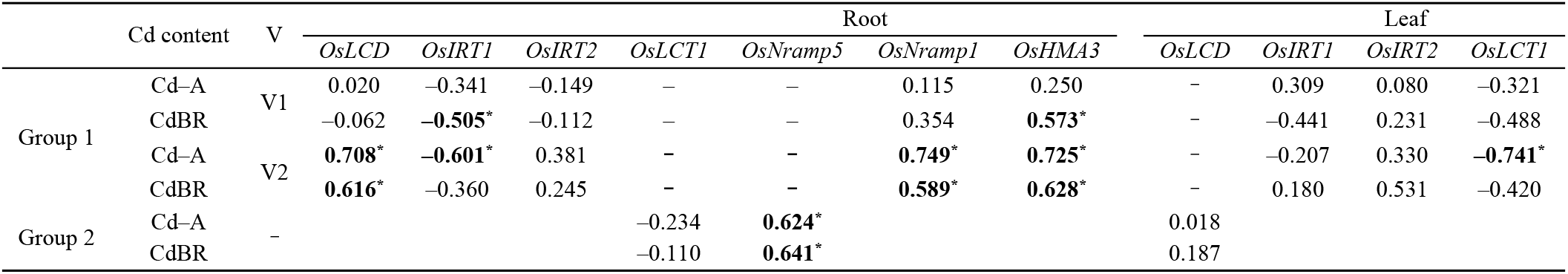
Correlation of Cd uptake transporter gene expression with Cd accumulation and brown rice Cd content in rice at maturity. Group 1 indicates expression of genes with varietal differences, and Group 2 with no varietal differences; Cd–A, Cd accumulation; CdBR, Cd content of brown rice.

**Table 8.**
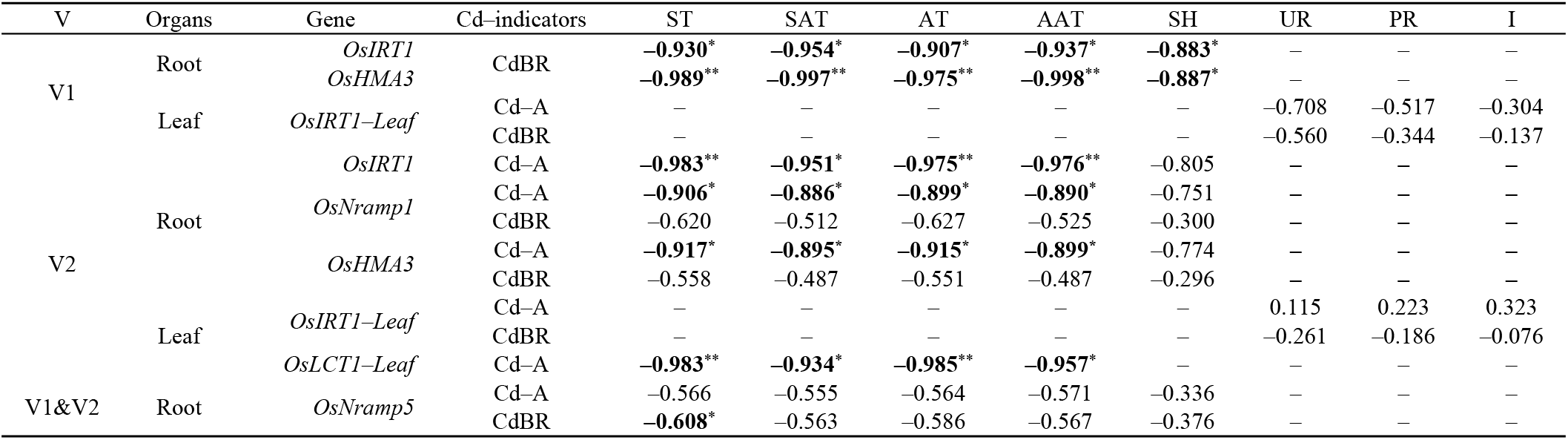
Partial relevancies of temperature and light factors with Cd-A and Cd-BR (with relative gene expression as a fixed factor)

## Disccusion

Cadmium pollution and the accumulation in rice, which then enters the human body through the food chain causes a potential threat to human health, is one of the major environmental problems all over the world[34]. Exploring the transport process of Cd in rice and constructing cultivation and management measures to reduce the absorption and accumulation of Cd in rice will help to improve rice growth and grain quality[3,35]. The absorption and distribution of Cd from soil to rice is a dynamic process, which relies on the absorption of other metal ion transporters through roots[36,37], root-to-shoot transport (xylem transport)[38,39] and source-sink transport (including seed loading) drive by phloem[40,41] to accomplish transportation and distribution among organs.

Since the climate e.g. precipitation, solar-radiation and air-temperature during growth period was different, the nutritional quality and yield potential were greatly different between early and late rice[42]. It was also proved that different water contents in paddy soils caused by different rainfall amount caused great variation in Cd accumulation in grains between early and later rice[26]. In this study, the mean number of days of field growth period (FGP) for the two varieties under different SDs was 83.0 and 83.2 days, with coefficients of variation of 0.76% and 1.68%, respectively (S1-sheet3), and shown the response of FGP to SDs was highly consistent between the two varieties (*p* = 0.787, independent samples t-test), indicating that the variation in each temperature (AT, ST, AAT, SAT) and light (UR, PR, SH, I) factor under different SDs was mainly due to the different position of FGP on the field trial timeline. Therefore, the different temperature and light conditions obtained by the SDs in present experiment were highly justified.

Temperature is considered to have a great correlation with cadmium absorption and accumulation in rice. Increasing temperature decreased the organic matter content rapidly and promoted metal availability and plant uptake[43]. It was believed that since the reduction in formation of iron plaque and decreased soil porewater pH, warming increased Cd accumulation in root to shoot, boosted Cd translocation from root to shoot, and influenced root morphology and increased leaf transpiration and boosted the xylem stream, subsequently significantly increased total uptake of Cd/Cu by rice[44]. In contrast to the present study, there was little difference in the AT during the FGP among different SDs, but the indexes related to temperature such as AAT, ST, AST etc., showed extremely significant negative correlations with CdBR. On the one hand, it was shown that temperature was one of more sensitive factors affecting the CdBR; On the other hand, under S6 treatment, the average temperatures of the two varieties at maturing phase were 18.66°C and 20.97°C, respectively, the seed setting rate and yield under these two treatments were the lowest in both varieties (S1-sheet5), which may suggest that the filling rate were affected seriously, could be considered as that the organic-material-flow containing Cd was distributed to fewer grains, resulting in the significant increase of CdBR under these treatments.

The response of CdBR to SDs was consistent between the two varieties (p=0.278), but the variation in CdBR was much higher in V2 than V1 under different SDs, indicating that the CdBR of V2 was more sensitive to the temperature and light environment. In addition, the CdBR of V1 was invariably greater than the national standard of 0.2 mg/kg at any SD, researchers treated it as a high Cd accumulating variety consequently, while V2 as a low Cd accumulating variety did not show low Cd content (<0.2 mg/kg) at any environment/SD, and it could be not constantly feasible to be used as an emergency variety for low cadmium early rice in Hunan province. To obtain grains with V2 below the national standard for CdBR, it would be more reasonable to use it as a single-season medium rice or/and early maturing late rice (transplanting period is about early July) in Hunan province.

The expression of seven genes in the root system and four genes in the leaves differed significantly under different SDs with different levels of responsiveness, which provides a theoretical possibility to regulate the expression of genes related to Cd uptake and transport through SD selection, and thus regulate the uptake and accumulation of Cd in rice, and the CdBR as well. For genes that did not differ in relative expression between rice varieties but differ significantly among SDs, such as *OsLCT1* and *OsNramp5* in roots and *OsLCD* in leaves, such regulation could be more easily achieved by SD, while the regulation of genes that differ not only among SDs but also between varieties may be another complicated story.

Studies on the molecular mechanisms of Cd uptake and transport in rice have confirmed that the expression of a number of genes were associated with the accumulation and distribution of Cd in rice plants, and/or the Cd content of the rice grain as well, e.g. iron-regulated transporter *OsIRT1* have been proved to play some roles in Cd uptake in rice[45]; *OsNramp1* and *OsNramp5* were major transporters contribute to Cd transport in rice[18]; *OsHMA3* as a P1B-type of ATPase affected root-to-shoot cadmium translocation in rice by mediating efflux into vacuoles[46]; *OsLCT1* regulated cadmium transport into rice grain[47]. In this study, the expression of these genes mentioned above were verified that were regulated by temperature and light factors in both varieties (Table 6.); In the same way that the expression of *OsIRT1, OsNramp1, OsNramp5*, and *OsHMA3* in root of V1 and/or V2, and OsLCT1 in leaf of V2, were significant correlated to CdBR and/or Cd-A (Table 7.), suggesting a definite relationship between temperature and light conditions and CdBR and Cd-A.

Partial correlation analysis is the process of removing the effect of the third variable when two variables are simultaneously correlated with a third variable, and analyzing only the degree of correlation between the other two variables, determined by the R-value of the correlation coefficient. Partial correlation analysis showed that the effect of temperature on the CdBR was the main factor in the different temperature and light conditions created by the SD settings. Limited the expression of *OsIRT1, OsNramp1*, and *OsNramp5* contributed to the reduction of cadmium content in brown rice[30,48], and the partial correlation between temperature and CdBR also indicated that increased temperature contributed to the reduction of Cd in V1 and/or V2. Nevertheless, in low Cd-accumulating cultivars, *OsHMA3* functions to sequester Cd to the root vacuoles, resulting in less Cd translocation from the roots to the shoots and grains, while in high Cd-accumulating cultivars, loss of function of OsHMA3 resulted in high root-to-shoot translocation of Cd[49]. The down-regulation of *OsMHA3* expression by increased temperature may have led to an increase CdBR in V 1and Cd -A in V2, but had no significant enhanced CdBR in V2, further confirmed that the increase in CdBR was mainly executed by the transport capacity of Cd in the phloem[50].

## Conclusion

Temperature and light conditions during rice FGP could be adjusted by SD. Different temperature and light conditions showed different expression levels of genes related to Cd uptake and transport in rice, thus affected Cd-A and CdBR. Increased in ST, SAT, AT and AAT down-regulated *OsIRT1, OsNramp1* and *OsNramp5* to reduce Cd-A and CdBR. In the double-season rice growing area of Hunan, Zhuliangyou 819 was grown as a single-season medium rice or early maturing late rice could limit the Cd content of brown rice to below the national safety standard level.

## Supplementary Materials

S1: sheet1, hourly temperature detected by the micro meteorological station (Vantage Pro 2, USA) installed in the field from 17 May to 22 October 2018; sheet2, daily average temperature; sheet3, major growth stages and their number of days for the two experimental varieties; sheet4, daily air and soil temperature and light factor indicators measured from 5 May to 20 October 2018; sheet5, theoretical yields of the two experimental varieties and their yield components under different SD treatments; sheet6, relative expression of seven Cd uptake and translocation-related genes in roots of two rice varieties under different SD treatments at maturity stage; sheet7, relative expression of four Cd uptake and translocation-related genes in leaves of two rice varieties under different SD treatments at maturity.

## Acknowledgements

Natural Science Foundation of Hunan Province (2019JJ40124) is thanked for the financial support; Hunan Rice Research Institute (HRRI) is thanked for providing the seeds. Author Contributions

## Author Contributions

Data curation, Huan Xiao, Qiaomao Chen and Yu Kuang; Formal analysis, Qiaomao Chen and Yu Kuang; Resources, Huan Xiao and Hejun Ao; Supervision, Xuehua Wang; Writing – original draft, Zhongwen Rang; Writing – review & editing, Zhongwen Rang, Zhenxie Yi, Xuehua Wang and Hejun Ao.

